# Visual signals suppress Alpha Power Increases & Frequency Decreases before and after a Mindfulness Meditation Intervention for Problem Gambling

**DOI:** 10.1101/359257

**Authors:** Kaili-Larissa Martin, Farah Jindani, Nigel Turner, Joseph FX DeSouza

## Abstract

The use of mindfulness meditation (MM) in the treatment of problem gambling (PG), has been used effectively for over five years. However, the neural mechanisms responsible for the improvements are unknown. The literature describes healthy individuals with an increase in alpha power and a decrease in alpha frequency after eight weeks of mindfulness meditation, but it is unknown if changes are similar amongst individuals with PG. Using resting-state electroencephalography (rsEEG), we measured the changes in alpha oscillations before and after an eight-week mindfulness meditation intervention (MMi) and a pre/ post-five-minute mindfulness meditation body scan (MMb). For people with PG, we observed an increase in alpha power and decreased alpha peak frequency after the MMi, while the inverse was true for the MMb. The most considerable alpha rhythm changes occurred in the frontal and temporal lobes, areas sensitive to reward and sensory processing in PG. Our observed changes may reflect theories that MMi for PG may improve attentional control as hypothesized by previous research in alpha oscillations and cue-reward processing.

## Introduction

Mindfulness meditation (MM) has been used in conjunction with cognitive behavioural therapy as a treatment for individuals with problem gambling (PG) (Chen et al., 2014). In healthy individuals, eight weeks of MM results in a decreased alpha frequency and an increase in alpha amplitude (Cahn & Polich, 2006). We measured the alpha oscillations of PG to determine if an eight-week mindfulness meditation intervention (MMi) results in similar alpha changes. The study we have conducted is the first to analyze resting-state electroencephalography (rsEEG) data for this treatment group (PG). A small body of rsEEG research discusses alpha amplitude and frequency amongst those with PG (Cahn & Polich, 2006; Guo et al., 2017). Missing from the literature are the changes in alpha oscillations after a mindfulness meditation (MM) for treatment of PG. This novel study will contribute to the understanding of the currently unknown neural mechanisms of MM and PG treatment.

The MMi is founded on mindfulness-based cognitive therapy protocols that have been used as one of the treatment options for mental health conditions such as depression, anxiety and problem gambling (Barnhofer et al., 2015; Chen et al., 2014). Individuals who have participated in an MMi for PG reported improved control over gambling, greater task-specific attention and reduced automatic thought patterns (Chen et al., 2014). Traditionally, an MMi consists of eight weekly sessions, however, mindfulness is also practiced with a five-minute mindfulness meditation body scan (MMb). The MMb is a tool taught and practiced as a component of the MMi. It was hypothesized that an individual could learn to control alpha oscillations with the MMb by purposefully attending to externally cued locations of the body (Kerr et al., 2013).

Changes in alpha oscillations are measured by shifts in alpha peak frequency and alpha amplitude. rsEEG measures alpha amplitude and frequency changes by recording summed electrical activity on the surface of the scalp. Neurological activity is measured in the unit of power, or microvolts squared *(μV2*). Microvolts are transformed from the time domain to the frequency domain through a calculation called the Fast Fourier Transform (Akin, 2002). The Fast Fourier Transform is an algorithm that samples a signal over a period of time and divides it into single sinusoidal oscillations at specific frequencies each with their own amplitude (Akin, 2002). The distance from one peak of a wave to the another is termed *amplitude (μV)*. Amplitude squared is termed *power (μV2)*. The rhythmic activity of waves in the frequency that range between 8-14 Hz is referred to as the alpha wave. The alpha wave contains the most power amongst delta, theta, alpha, beta, and gamma identified in the brain (Kilmesch, 2012). The alpha rhythm has been found to be a marker of spontaneous cell firing which occurs during inactivity (Adrian & Matthews, 2010). The alpha rhythm disappears when the eyes are opened and during concentrated attention when the eyes are closed (Adrian & Matthews, 2010).

As alpha amplitude increases, weak cells are inhibited from firing, and blood flow decreases in the cortical area. As alpha amplitude increases, task-irrelevant stimuli from the external environment is filtered from cognitive processing (Klimesch, 2012). As cortical activation decreases, a selection of attentional resources occurs, and task-irrelevant information from the external environment is filtered. Alternatively, alpha amplitude decrease occurs as cell inhibition is released and cortical activity increases (Klimesch, 2012). With increased cortical activity, inhibition of sensory stimuli is difficult and reflected in decreased alpha amplitude (Klimesch, 2012).

The MMb may be used as a tool to control the regulation of thalamocortical activity by purposefully attending to parts of the body (Kerr et al., 2013). Alpha oscillations reflect changes in thalamic-cortical activity (Klimesch, 1999). The thalamus acts as a relay centre sending sensory input to different parts of the cortex. By controlling the regulation of thalamocortical activation loops, one may be able to modulate the transfer of sensory input received from the environment.

Changes in alpha frequency sub-bands reflect different cognitive processes, arousal (11 to 14 Hz), attention (9 to 10 Hz) and memories (7 to 8 Hz) (Klimesch, 2012; see Appendix). An increase in alpha frequency is associated with improved cognitive performance and memory (Klimesch, 1999). Alpha frequency is higher in childhood to early adulthood and decreases with aging (Chiang, 2011). Low alpha frequencies have been noted in individuals with low IQ and/or neurodegenerative diseases (Duffy et al., 1984; Nimmrich et al., 2015). More recently, cortical alpha has been shown to predict the confidence in an impending action (Kubanek et al., 2015) which could reflect motor preparation of an impending action (DeSouza et al., 2003) or motor efference copy signals (DeSouza et al., 2000; DeSouza et al., 2002) which also employ thalamocortical loops.

PG has displayed lower alpha amplitude in the left frontal (F7) and temporal (T7) lobe and activation of reward centres during high-risk decision making (Miedl et al., 2014). MM has resulted in an alpha power increase in the frontal and temporal lobes in healthy individuals (Cahn & Polich, 2006). MM has also resulted in a reduced neurological response to rewards (Tang et al., 2009). PG exhibits a dysregulation in neural networks involved in gambling-related cue and reward responses (Murch & Clark, 2015). External stimuli from gambling such as lights or noise produce physiological sensations. The insula is sensitive to physiological signals and encodes an emotion via the amygdala to be later recalled by the hippocampus (Davis & Whalen, 2001; Gupta et al., 2011). PG has shown to create a heightened sensitivity to triggering cues, previously associated with gambling, resulting in physiological arousal. Conditioned gambling cues activate the mesolimbic system, resulting in cravings for gambling-related rewards (Tang et al., 2015). Attentional bias towards gambling cues occurs with PG (Janssen et al., 2015; Murch & Clark, 2015). Greater internalized control over attention to gambling-related cues may be beneficial in the prevention of PG behaviours. It has been hypothesized MMi compliments treatment for PG because it leads to greater attentional control over internal and external stimuli (Chen et al., 2014). Internalized control over attention to external stimuli is also positively correlated with alpha oscillations (Klimesch, 2012) and thus our examination of the neural mechanisms in this report.

rsEEG data was collected in two separate conditions, eye’s open (EO) and eye’s closed (EC) for 6-minutes. The EO and EC conditions reflect different measures of baseline alpha activity (Barry et al., 2007; Adrian & Matthews, 2010). In the EO condition, visual stimulation leads to information processing and global activation of the cortex (VaezMousavi et al., 2007). The EO condition is a better measure of the resting state baseline while visual processing takes place before a task-specific situation (VaezMousavi et al., 2007). The EC condition measures arousal or one’s energetic state at a given moment (VaezMousavi et al., 2007) and would use differing corticothalamic loops than with EC. The EC condition putatively leads to a decrease in cortical activation and alpha waves are more dominant (Barry et al., 2007). rsEEG data was collected and analyzed in both conditions (EO/EC) to compare and contrast different aspects of the dynamics of alpha oscillations. Alpha frequency should be higher in the EO condition than the EC condition due to the visual processing and heightened cortical activity.

## Methods

rsEEG recordings used an Epoc Emotiv 14 channel headset with TestBench, the accompanying software that displays and records rsEEG data. Saline solution was applied to the felt tips of the EEG headset, the participant’s head was measured, and the headset was placed on the head in accordance with electrode measurement positions: AF3, AF4, F3, F4, F7, F8, FC5, FC6, P7, P8, T7, T8, O1, O2 and reference electrodes at P3/P4, based on the modified combinatorial nomenclature system (Adjouadi et al., 2004). Sensors were adjusted to ensure the best impedance. Participants were instructed to look at a *cross* displayed on a laptop screen in front of them via MediaLab software (v2012). A three-minute recording collected rsEEG data once with the participant’s eyes open (EO) and once with the participant’s eyes closed (EC). Data markers were set to indicate the beginning and end of the EO and EC conditions, sent by Virtual Serial Port Driver (Version 7.1, Eltima Software, 2013, Bellevue, WA) to TestBench and saved as an EDF file (Di Nota et al., 2017).

A total of 28 self-selected volunteers with PG took part in this study. Compensation of $40 was provided to each participant before and after the MMi. Participants completed a mindfulness-based body scan (MMb). The MMb was led by a clinician or researcher with more than ten years of MM practice. The participants were then briefed on the MMb, and the lights in the room were dimmed. Participants were invited to sit in a chair and close their eyes if it suited their comfort. Next, participants were instructed to bring their attention to each part of their body from toe to head beginning with the left or right side, chosen by the instructor. Participants were asked to direct attention to the physical body or their breath upon awareness of internal thought dialogue. Upon completion of the five-minute MMb, the lights were turned on, and rsEEG data was collected for the second time (t2). Participants then took part in an eight-week MMi led by a clinician at the Centre for Addiction and Mental Health (CAMH) services in Toronto, Ontario. The MMi was led in accordance with mindfulness-based cognitive therapy and mindfulness-based relapse prevention protocols. Post MMi, participants returned to CAMH for data collection. Participants first completed the STROOP test, followed by rsEEG data collection (t3), an MMb, then a fourth data collection point (t4).

### Data Preprocessing

Six minutes of rsEEG data were preprocessed into two-second epochs using the Fieldtrip toolbox (Oostenvelde et al., 2011) in Matlab v7.12.0, R2011a (The Mathworks, Inc., Natick, MA). An independent component analysis on the Fourier transform of EEG signal was used to reject artifacts. Components were visually inspected before rejection. Artifacts were removed, due to putative eye blinks, muscular movements and signal noise. Power spectrum data was transformed using a fast Fourier transformation (for data pipeline see https://bitbucket.org/joelab/eeg-tutorial/wiki/Home).

### Data Analysis

Alpha power was log-transformed for statistical normalization. Data was then imported to SPSS v24 for analysis. A linear mixed model was used to accommodate for missing cells of data across the sample size (n=28) and random effects (Aarts et al., 2014; Wang & Goonewardene, 2004). Time was treated as a continuous variable, and subject variables were fixed. The model assigned each participant their own point on the y-intercept which randomly deviates from the group mean at each time interval along the x-axis. The factor analytic, first order covariance structure was used to accommodate unequally spaced time intervals with maximum likelihood (ML) estimation. Histograms were created, and normality was assessed. Model fit was assessed with Akaike information criterion (Akaike, 2011). Post-hoc mean comparisons were made using the Bonferroni adjustment.

The model had a total of eight repeated measures observations per participant (4-time intervals x 2 EO/EC conditions). Level one consisted of participants (N=28). In level two, each participant had two pairs of data collection pre- and post MMi, creating unequally spaced time intervals. Time point one (t1) consisted of rsEEG data collection, followed by a five-minute MMb at time point two (t2). A second rsEEG measurement was taken immediately after. Time point three (t3) and time point four (t4) consisted of the same pattern of data collection after the MMi.

Missing data (MAR) was <5% of cases. An unbalanced data set was due to participant drop out, failure of EEG marker transfer to TestBench, unresolved errors and muscular artifacts or noise creating unusable rsEEG data. With unbalanced data sets, estimates of fixed effects are based on the maximum and minimum likelihood methods (ML) (Field, 2013). The ML estimation of variance was selected for its robustness as compared to the restricted maximum likelihood method (REML) when looking at fixed effects (Field, 2013). We hypothesized that the group of participants, the fixed effect, would show a change in alpha oscillations over time, therefore use of the ML was appropriate.

## Results

Electrodes in the left and right frontal and temporal areas changed after the MMi and MMb (see Figure 1). We hypothesized an increase in alpha power and decrease in alpha frequency after the MMi and MMb. After the MMi, a significant increase in alpha power and a decrease in frequency was observed, however, after the MMb, the inverse was observed (see Figure 2).

**Figure 1.**
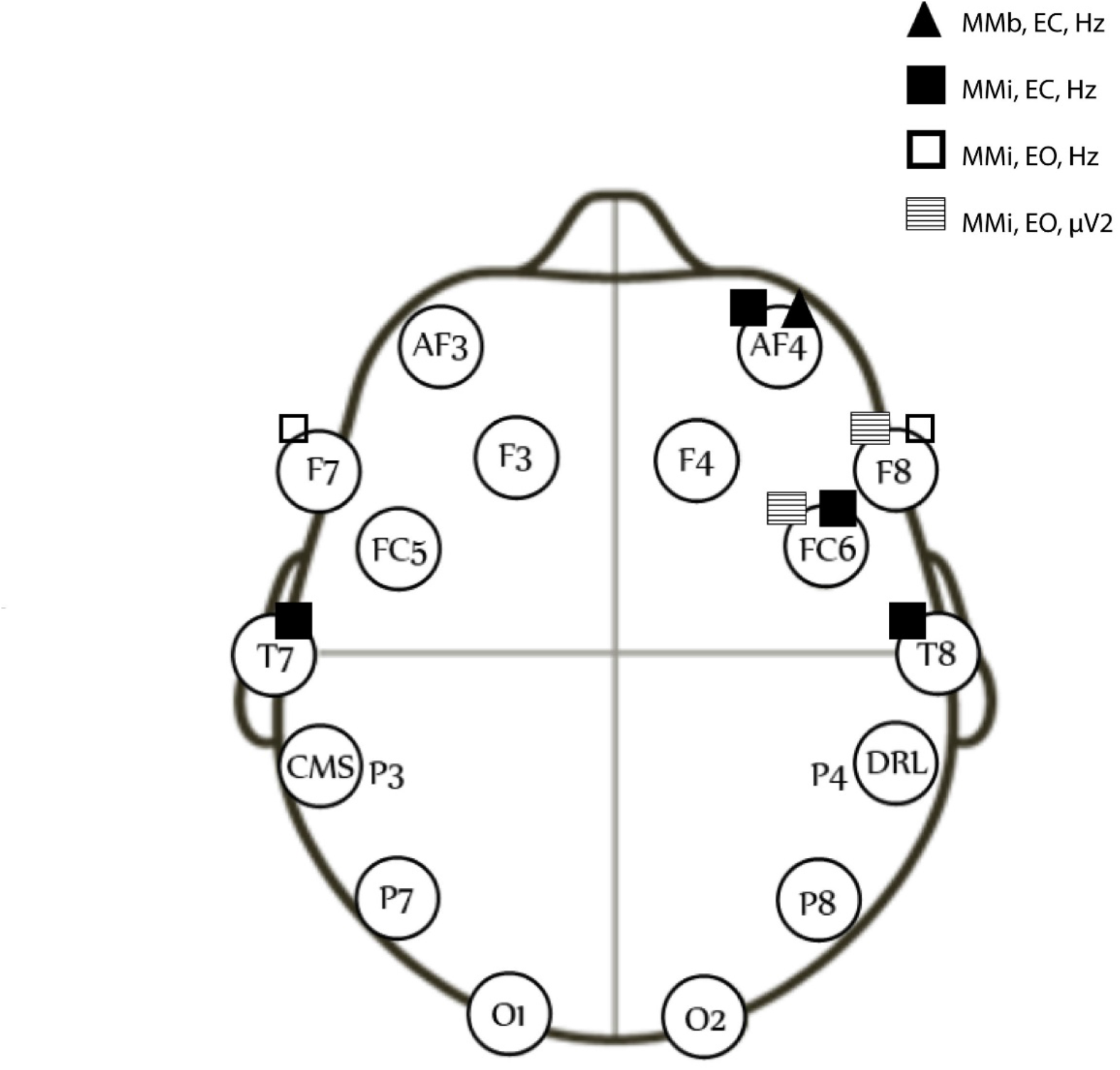
Electrodes with significant changes in alpha amplitude ((µV2) and alpha peak frequency (Hz) after the MMi and MMb in both the EC and EO condition. Adapted from Torres-García, A. A., Reyes-García, C. A., Villasenor-Pineda, L., & Ramírez-Cortés, J. M. (2013). Análisis de senales electroencefalográficas para la clasificaciÓn de habla imaginada. Revista mexicana de ingeniería biomédica, 34(1), 23-39.

**Figure 2.**
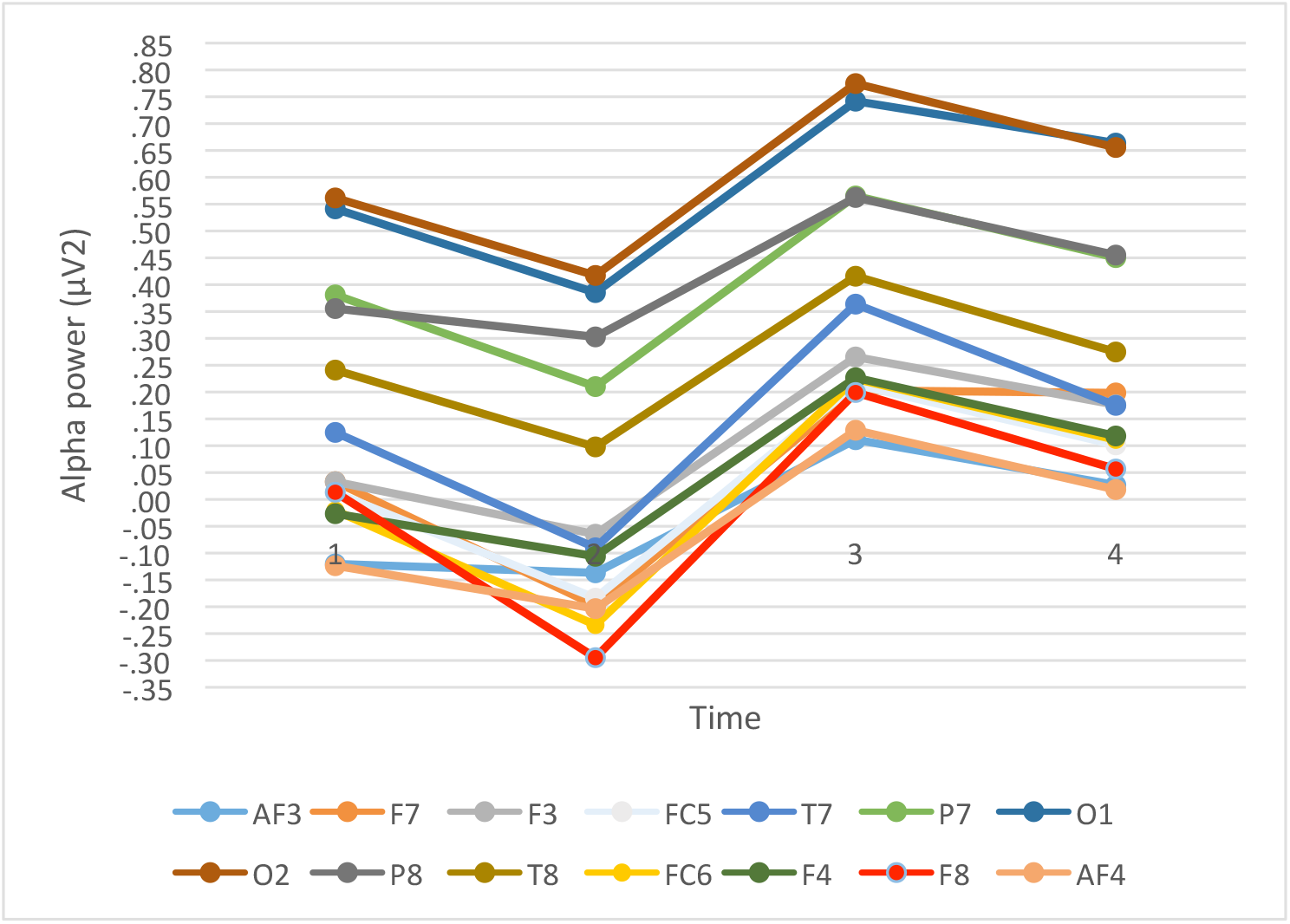
Alpha amplitude decrease in the EO condition after the MMb (t1 to t2 and t3 to t4) and alpha amplitude increase after the MMi (t1 to t3 and t2 to t4).

### Log alpha power (µV2)

Looking at the mean group scores of each of the 14 electrodes in the EO condition, a significant change over time was found F (3,18.51) = 71.00, p<.001. A significant increase of 0.14 units between t1 and t3 p<.001, an increase of 0.13 units between t1 and t4 p<.001 an increase of 0.20 units between t2 and t3, p<.001, and an increase of 0.20 between t2 and t4, p<.001.

Looking at the 14 electrodes as a group in the EC condition, we found a significant change over time, F (3,19.32) = 74.41, p<.001. A significant increase of 0.14 between t2 and t3, 0.15 between t1 and t3, .07 between t1 and t4, and a decrease of .09 between t3 and t4. All correlations are statistically significant for the EO condition with p<.001 and range from 0.721 to 0.951. All correlations are statistically significant for the EC condition with p<.001 and range from 0.795 to 0.955.

With subject scores fixed and electrodes individually modeled as the dependent variable, the overall trend of alpha amplitude decrease after the MMb and alpha amplitude increase after the MMi in the EC condition is displayed in Figure 3. Results displayed the same trend in the EO condition.

**Figure 3.**
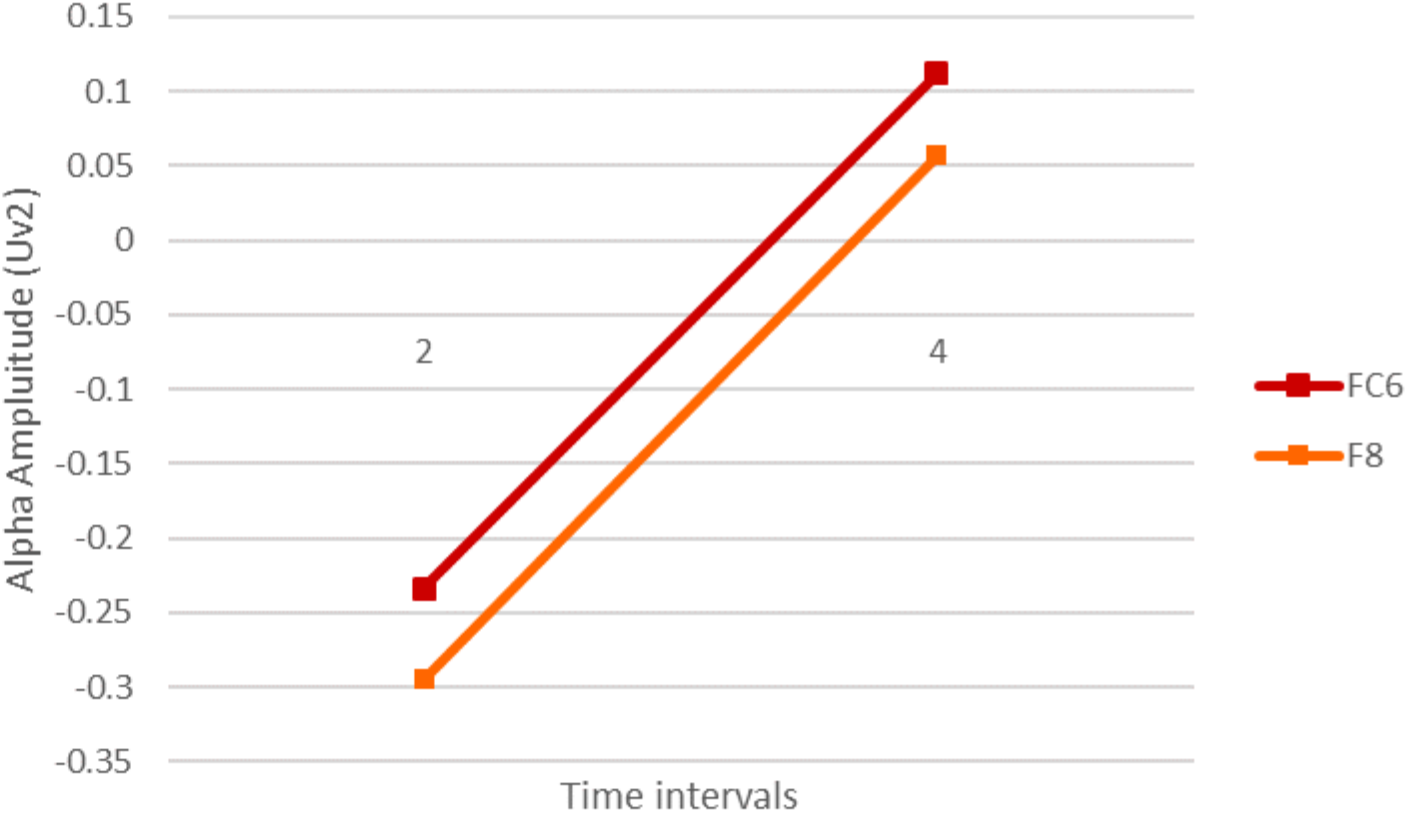
Alpha amplitude increase in the right frontal lobe post MMi. F8 showed a significant change in alpha amplitude, F (3,30.616) = 4.80, p=.007, an increase, b=0.538 (95% CI 0.131-0.946) SE=0.14 from t2 to t4, p=.005. FC6 showed a significant change, F (3,14.850) = 4.353, p=.022, with an increase b=0.287 (95% CI 0.026-0.547) SE=0.14 from t2 to t4, p=.025.

With subject scores fixed, and electrodes FC6 and F8 individually modeled as the dependent variable, alpha power increased significantly after the MMi. (See Figure 2). In FC6 and F8 alpha power increased significantly in the right frontal lobe post-MMi. F8 showed a significant change in alpha amplitude, F (3,30.616) = 4.80, p=.007, an increase, b=0.538 (95% CI 0.131-0.946) SE=0.14 from t2 to t4, p=.005. FC6 showed a significant change, F (3,14.850) = 4.353, p=.022, with an increase b=0.287 (95% CI 0.026-0.547) SE=0.14 from time point 2 to 4, p=.025.

## Alpha Frequency (Hz)

With individual electrodes modeled as the dependent variable, a significant decrease in alpha frequency was found in both the left and right frontal and temporal regions after the MMi as displayed in Table 1. In the EO condition, the left frontal electrode (F7) and the right frontal, temporal region (F8) displayed a significant decrease in alpha frequency after the MMi as seen in Figure 4.

**Figure 4.**
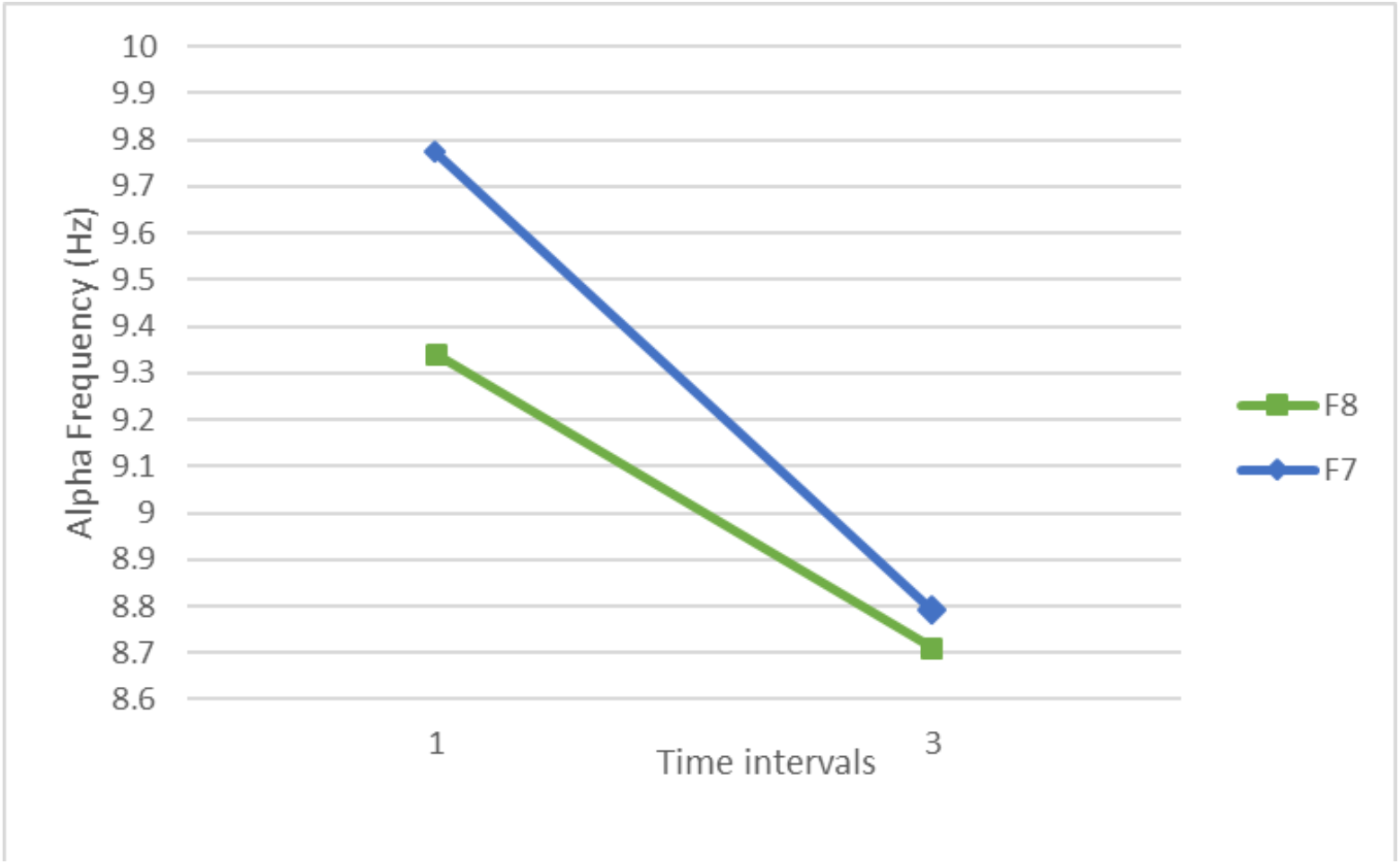
Alpha frequency decrease in the left and right frontal electrodes. Electrode F7 displayed a significant decrease after the MMi, b=0.820 (95% CI 1.296-0.345) SE=0.169, from t1 to t3, p=<.001 and from t1 to t4, b=0.855 (95% CI 0.144-1.565) SE=0.250, p=<.001, after the MMi. A significant change was also found in the right frontal, temporal region (F8). There is a significant decrease after the MMi b=0.677 (95% CI 0.050-1.304) SE=0.218 from t1 to t3, p=.029.

**Table 1.**
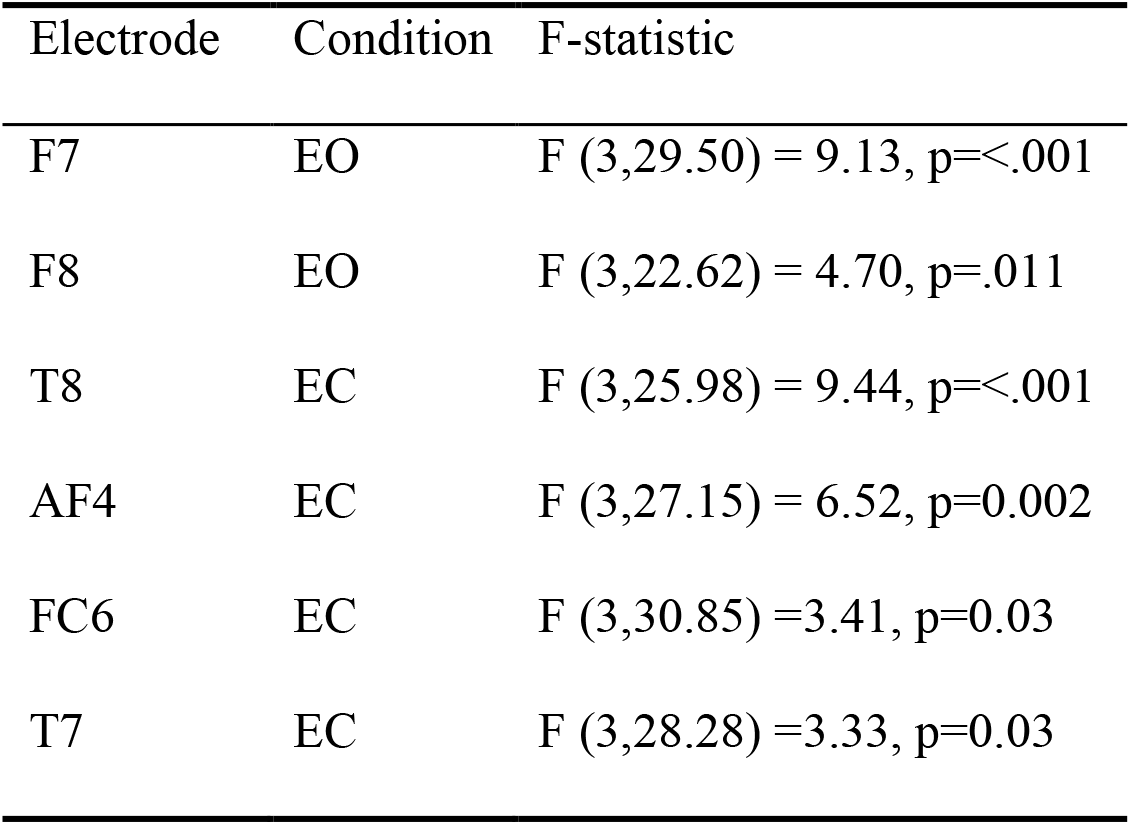
Electrodes with significant alpha frequency changes in the EO and EC conditions.

| Electrode | Condition | F-statistic |
| --- | --- | --- |
| F7 | EO | $F(3,29.50) = 9.13, p < .001$ |
| F8 | EO | $F(3,22.62) = 4.70, p = .011$ |
| T8 | EC | $F(3,25.98) = 9.44, p < .001$ |
| AF4 | EC | $F(3,27.15) = 6.52, p = 0.002$ |
| FC6 | EC | $F(3,30.85) = 3.41, p = 0.03$ |
| T7 | EC | $F(3,28.28) = 3.33, p = 0.03$ |

In the EC condition, electrodes T7 and T8 showed a significant decrease in alpha frequency after the MMi. (Figure 5).

**Figure 5.**
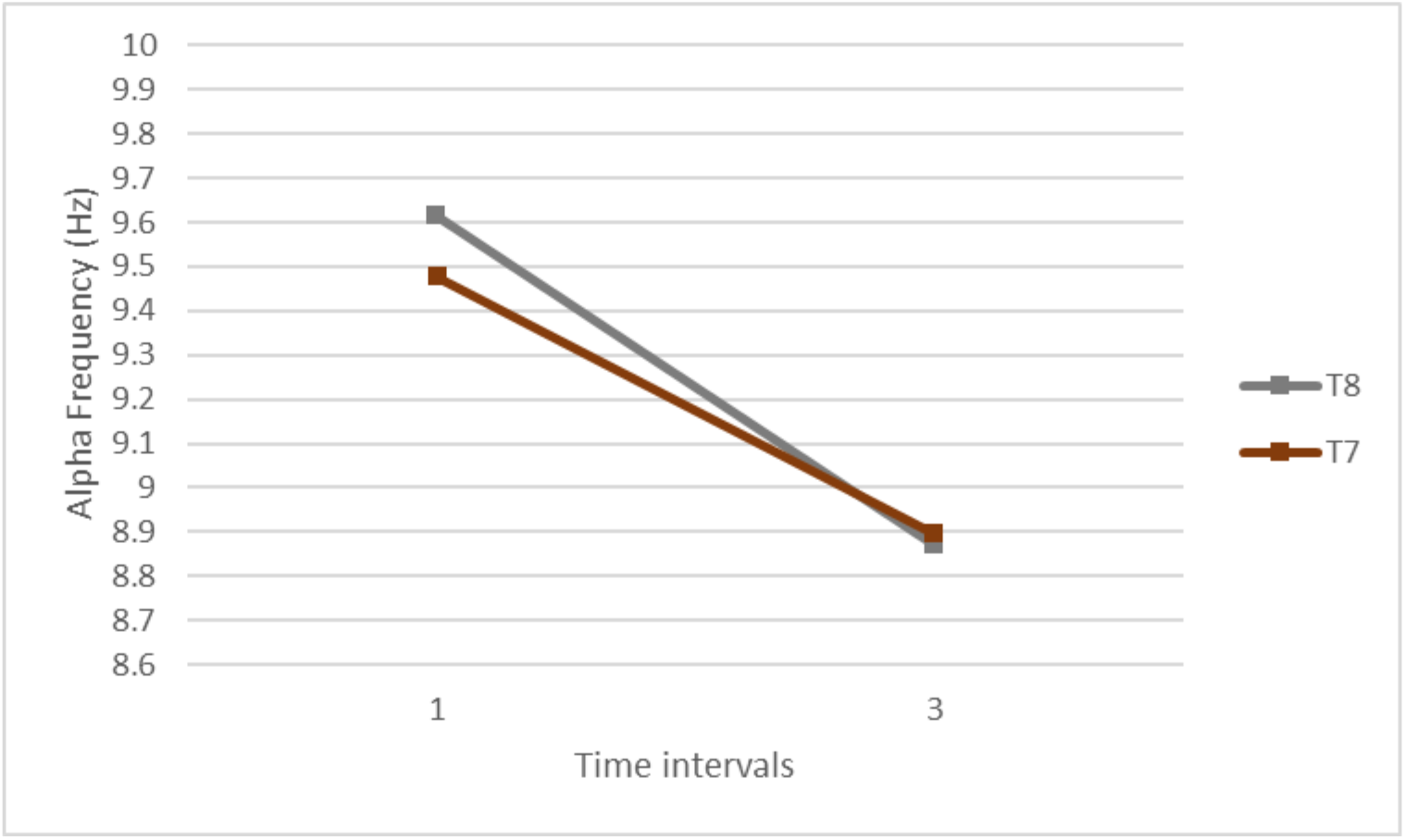
Electrodes T8 and T7 showed a significant decrease after the MMi. T8 showed a significant decrease b=0.005 (95% CI 0.256-1.670) SE=0.238 from t1 to t3, p=.005. There is also a significant decrease b=0.002 (95% CI 0.283-1.479) SE=0.240 from t1 to t4, p=.002. Electrode T7 showed a significant effect of time, with no significant post hoc effects.

**Figure 6.**
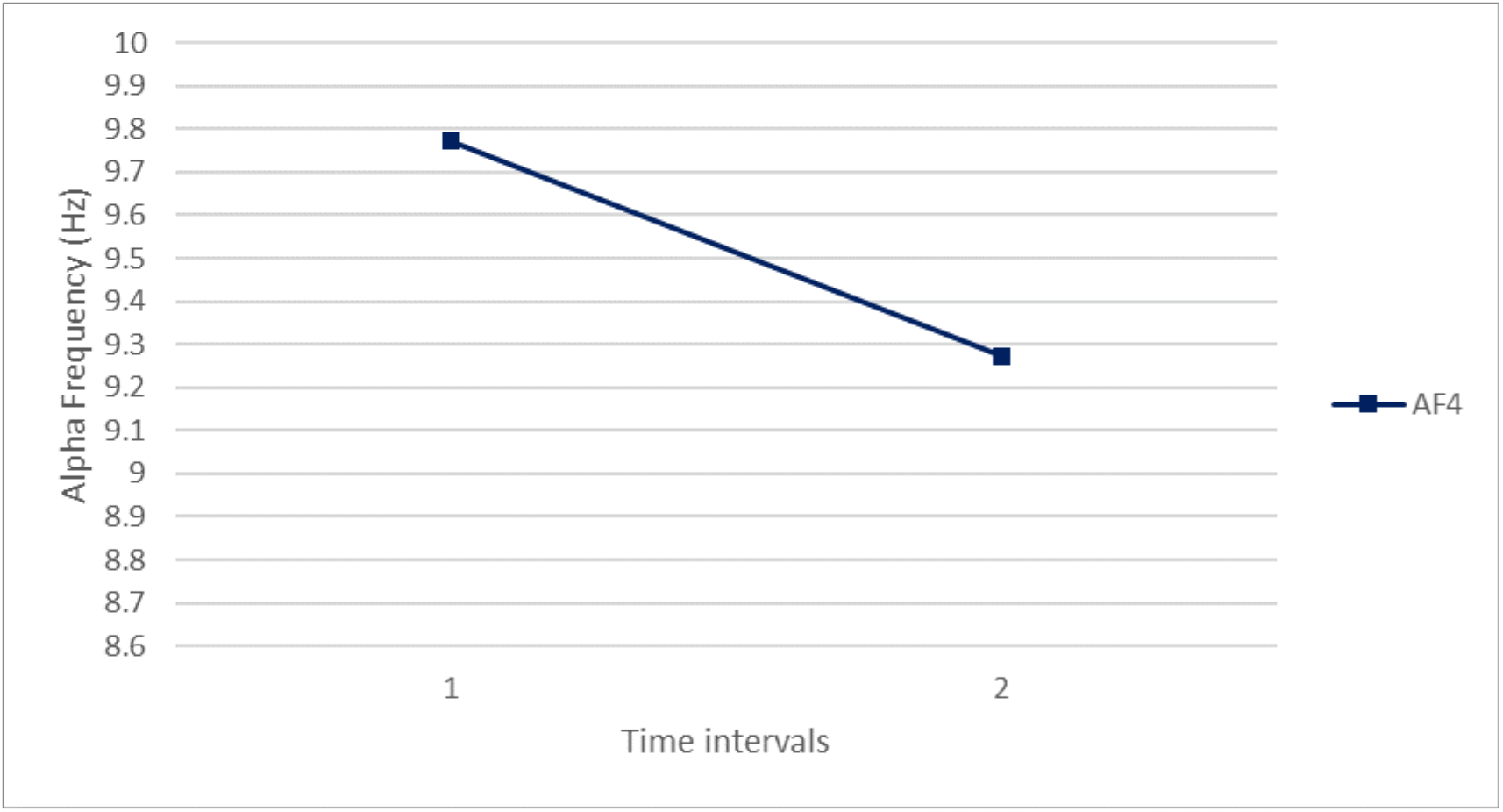
AF4 showed a significant decrease after the MMb. F(3,27.153)6.52,p=0.002 b=0.465 (95% CI 0.810-0.120) SE=0.119 from t1 to t2, p=.005.

FC6 showed a significant effect of time (see table 1), with no significant effect in post hoc comparisons. AF4 showed a significant decrease b=0.465 (95% CI 0.120-0.810) SE=0.119 from t1 to t2, p=.005. There was also a significant increase b=0.388 (95% CI 0.060-0.716) SE=0.119 from time point 2 to time point 3, p=.013 (See figure 4).

## Discussion

We hypothesized an increase in alpha power and decrease in alpha frequency after the MMi and MMb in the frontal and temporal lobes in people with PG. The overall trend after the MMi of an increase in alpha power and a decrease in frequency was observed, however, the inverse was true after the MMb. Individual electrodes in the fronto-central and temporal regions changed significantly as predicted based on previous research in normal people undergoing mindfulness meditation.

### Mindfulness meditation intervention - MMi

#### Frequency (Hz)

After the MMi, a decrease in frequency was observed in the left and right frontal lobes (F7 and F8) in the EO condition. The EO condition represents visual processing before a task-specific situation with heightened activation of the entire cortex (Barry et al., 2007). Our results suggest a reduction in cortical activity in the frontal lobes during visual processing before a task-specific situation. A decrease in frequency was also found in the left and right temporal lobes (T7 and T8) in the EC condition. The EC condition represents internal arousal or energetic state at any given moment (VaezMousavi et al., 2007).

Reduced frequency in the temporal lobes may represent a reduction in sensory information transfer at any given moment. Previous research has hypothesized that theT8 represents a gating mechanism for incoming sensory information to be relayed to the cortex (Jensen & Mazaheri 2010; Klimesch, 2007). Changes in alpha frequency reflect attentional resources and the transfer of sensory information to the cortex (Klimesch, 2012).

After the MMi, a significant decrease in alpha frequency was found in the left frontal and temporal areas (F7 and T7) in the EO and EC conditions respectively. Previous studies have shown these areas to be overactive with a low alpha amplitude during risky decision making (Miedl et al., 2014). The decrease in alpha frequency may result in lower cortical activation of areas related to high-risk decision making. A lower frequency in F7 and T7 may represent less information transfer between the sensory system and the cortex. iwPG have a strong reaction to gambling cues attended to by the sensory system and hypersensitivity of cortical areas involved in risk and reward. MMi may benefit iwPG when presented with a gambling cue by improved attentional control over task-irrelevant sensory stimuli.

Frequency decrease was also found amongst electrodes and FC6 and AF4, the right fronto-central and front-most electrodes. These results coincide with changes found amongst healthy individuals after eight weeks of MM. The asymmetric changes warrant further research on iwPG.

#### Log alpha power (µV2)

F8 and FC6 increased in amplitude after the MMi. Previous studies have found alpha amplitude to be positively correlated with internalized control over attendance to the task-irrelevant sensory processing of environmental stimuli. Greater attentional control would be beneficial for iwPG who show attentional bias to gambling-related cues in the environment.

### Mindfulness meditation body scan - MMb

#### Alpha Frequency (Hz)

The trend of alpha frequency increase was observed after the MMb in both the EC and EO conditions. However, a significant decrease in frequency in electrode AF3 was identified. A significant effect of time was observed in FC6, but post hoc comparisons failed to identify the time interval. This may have been due to our very conservative criteria of multiple comparisons employing the Bonferonni correction. FC6 increases after the MMb, however, the significance of the time period is unknown. As a trend, more electrodes saw an increase in frequency after the MMb than not. Significant changes in electrodes FC6 and AF4 with significant post hoc comparisons in AF4 result in the observation in the right frontal lobes changes in frequency over time, with a decrease in AF4 after the MMb. Alpha peak frequency has been hypothesized to represent the transfer of sensory information from the thalamus to the cortex (Klimesch, 1999). Our results may represent the cognitive processing that occurs during the MMb as one purposefully directs attention to each body part across time at the direction of the instructor (authors FJ and K-LM). Usually, the thalamus inhibits the senses of the external body until required, so the MMb requires increased attentional demands and cognitive processing upon direction from the instructor. It was hypothesized that the MMb might be used as a tool to learn to control alpha oscillations, by regulation of communication between the thalamus and the cortex (Kerr et, al. 2013). Over time the use of this tool may provide greater control over attendance to or suppression of irrelevant sensory input that is required in addiction research in our population and could also be applied to others.

#### Log-alpha power (µV2)

A decrease in alpha amplitude was observed after the MMb. A decrease in amplitude putatively results in released inhibition of neuronal firing, increased cortical activity and cognitive performance. The improved cognitive performance may be beneficial for iwPG who display gambling related thinking distortions that have been shown to decrease with rational thought (Emond & Marmurek, 2010). For example, if one is tempted to gamble, a five-minute body scan phone app could be a useful tool to improve cognitive clarity and increase rational thinking as opposed to automatic thinking.

Alpha power decrease may have also occurred as participants think about body movement. For example, as attention is directed from one body part to another, or the practitioner suggests subject’s direct attention to the chest as it moves in and out while breathing. Alpha power decrease has been found when one cognitively prepares for movement (Brinkman et al., 2014).

All observed changes in alpha oscillations that occurred in the subbands may be related to attention (9 to 10 Hz) and memory (7-8 Hz). We plan to conduct follow-up studies to analyze working memory and attention in more detail.

Limitations of our initial preliminary study include the lack of a control group; however, future research is planned in the next stage. The changes in alpha amplitude and frequency, amongst iwPG after an MMi, are not yet found in the literature. The results assist in understanding neural changes after MM practices in iwPG. Future research may investigate whether MMi for iwPG could lead to better internalized control over cues that lead to problem gambling actions (Kubanek et al., 2015). It has been hypothesized that PG occurs with a strong cue-related craving during high-risk decision making (Miedl et al., 2014). iwPG show attentional bias to gambling-related cues relayed to the cortex by the sensory system (Tang et al., 2015). The MMi may provide protection from PG by unconsciously reducing attendance to environmental cues that trigger cravings and behaviour. The MMb may be a tool one can consciously implement to improve problematic gambling behaviours.

## Acknowledgements

We thank Peter Chen for allowing this research in his MMi class and having JFXD in his first cohort class. K-LM wrote this preprint report as part of her honours thesis began in 2015-16. FJ supervised K-LM onsite. NT and JFXD provided funds from OPGRC. JFXD edited versions of this manuscript. All authors approve of the current version for submission.

## Appendix

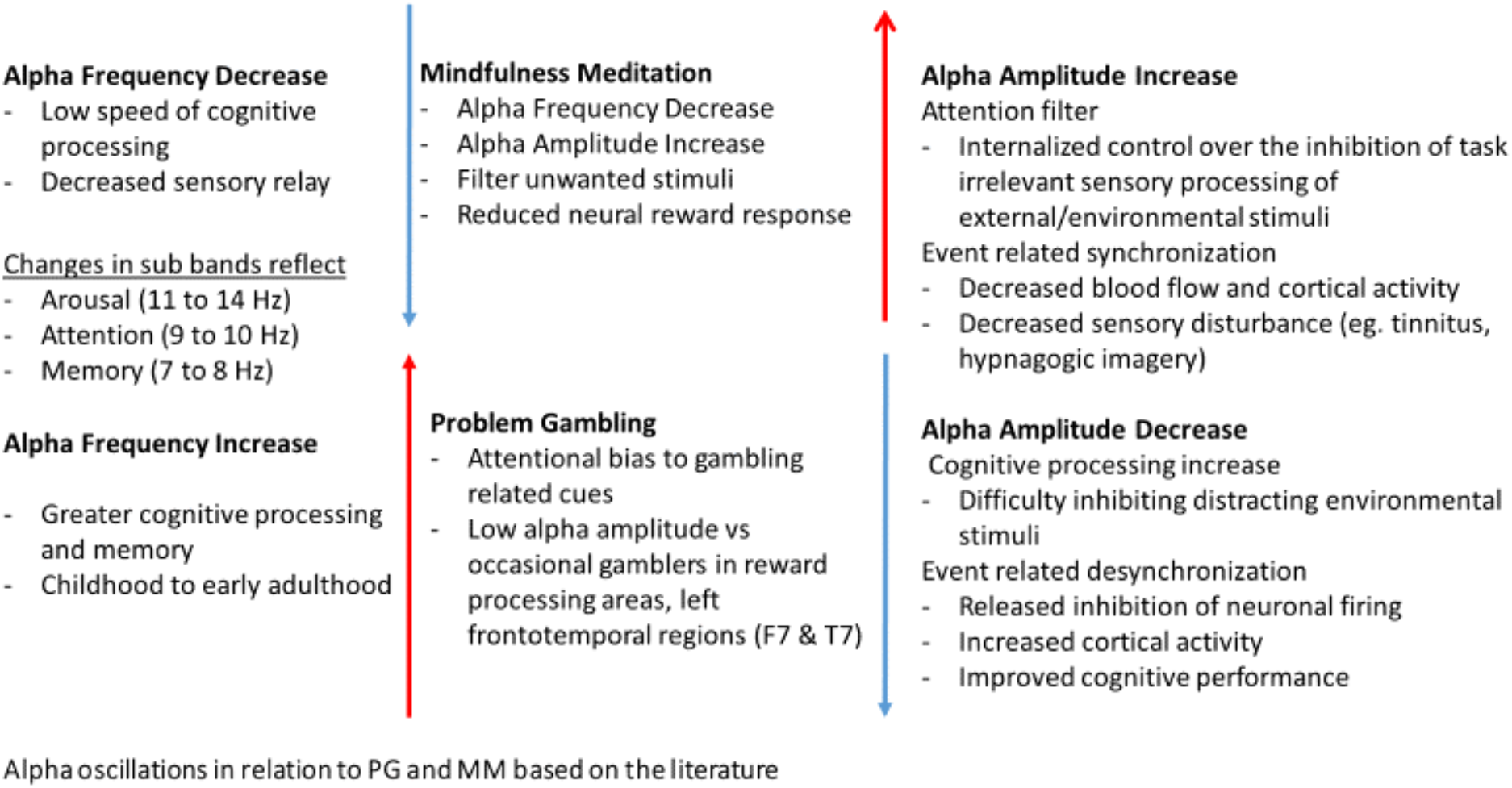

## References

1. Aarts, E., Verhage, M., Veenvliet, J. V., Dolan, C. V., & van der Sluis, S. (2014). A solution to dependency: using multilevel analysis to accommodate nested data. Nature neuroscience, 17(4), 491–496.

2. Adrian, E. D., & Matthews, B. H. (2010). The Berger rhythm: potential changes from the occipital lobes in man. Brain, 57(4), 355–385.

3. Akaike, H. (2011). Akaike’s information criterion. In International encyclopedia of statistical science (pp. 25-25). Springer Berlin Heidelberg.

4. Barnhofer, T., Huntenburg, J. M., Lifshitz, M., Wild, J., Antonova, E., & Margulies, D. S. (2015). How Mindfulness Training May Help to Reduce Vulnerability for Recurrent Depression A Neuroscientific Perspective. Clinical Psychological Science, 2167702615595036.

5. Barry, R. J., Clarke, A. R., Johnstone, S. J., Magee, C. A., & Rushby, J. A. (2007). EEG differences between eyes-closed and eyes-open resting conditions. Clinical Neurophysiology, 118(12), 2765–2773.

6. Brinkman, L., Stolk, A., Dijkerman, H. C., de Lange, F. P., & Toni, I. (2014). Distinct roles for alpha-and beta-band oscillations during mental simulation of goal-directed actions. Journal of Neuroscience, 34(44), 14783–14792

7. Cahn, B. R., & Polich, J. (2006). Meditation states and traits: EEG, ERP, and neuroimaging studies. Psychological bulletin, 132(2), 180.

8. Chen, P., Jindani, F., Perry, J., & Turner, N. L. (2014). Mindfulness and problem gambling treatment. Asian Journal of Gambling Issues and Public Health, 4(1), 1.

9. Davis, M., & Whalen, P. J. (2001). The amygdala: vigilance and emotion. Molecular psychiatry, 6(1), 13.

10. DeSouza, J. F., Dukelow, S. P., Gati, J. S., Menon, R. S., Andersen, R. A., & Vilis, T. (2000). Eye position signal modulates a human parietal pointing region during memory-guided movements. Journal of Neuroscience, 20(15), 5835–5840.

11. DeSouza, J. F., Dukelow, S. P., & Vilis, T. (2002). Eye position signals modulate early dorsal and ventral visual areas. Cerebral Cortex, 12(9), 991–997.

12. DeSouza, J. F., Menon, R. S., & Everling, S. (2003). Preparatory set associated with pro-saccades and anti-saccades in humans investigated with event-related FMRI. Journal of Neurophysiology, 89(2), 1016–1023.

13. Di Nota, P. M., Chartrand, J. M., Levkov, G. R., Montefusco-Siegmund, R., & DeSouza, J. F. (2017). Experience-dependent modulation of alpha and beta during action observation and motor imagery. BMC neuroscience, 18(1), 28.

14. Duffy, F. H., Albert, M. S., McAnulty, G., & Garvey, A. J. (1984). Age-related differences in brain electrical activity of healthy participants. Annals of neurology, 16(4), 430–438.

15. Emond, M. S., & Marmurek, H. H. (2010). Gambling related cognitions mediate the association between thinking style and problem gambling severity. Journal of Gambling Studies, 26(2), 257–267.

16. Field, A. (2013). Discovering statistics using IBM SPSS statistics. Sage.

17. Gupta, R., Koscik, T. R., Bechara, A., & Tranel, D. (2011). The amygdala and decision-making. Neuropsychologia, 49(4), 760–766.

18. Janssen, L. K., Sescousse, G., Hashemi, M. M., Timmer, M. H. M., ter Huurne, N. P., Geurts, D. E. M., & Cools, R. (2015). Abnormal modulation of reward versus punishment learning by a dopamine D2-receptor antagonist in pathological gamblers. Psychopharmacology, 232(18), 3345–3353.

19. Kerr, C. E., Sacchet, M. D., Lazar, S. W., Moore, C. I., & Jones, S. R. (2013). Mindfulness starts with the body: somatosensory attention and top-down modulation of cortical alpha rhythms in mindfulness meditation. Frontiers in human neuroscience, 7, 12.

20. Klimesch, W., Sauseng, P., & Hanslmayr, S. (2007). EEG alpha oscillations: the inhibition– timing hypothesis. Brain research reviews, 53(1), 63–88.

21. Klimesch, W. (2012). Alpha-band oscillations, attention, and controlled access to stored information. Trends in cognitive sciences, 16(12), 606–617.

22. Kubanek, J., Hill, J., Snyder, L. H., & Schalk, G. (2015). Cortical alpha activity predicts the confidence in an impending action. Frontiers in neuroscience, 9, 243.

23. Miedl, S. F., Fehr, T., Herrmann, M., & Meyer, G. (2014). Risk assessment and reward processing in problem gambling investigated by event-related potentials and fMRI-constrained source analysis. BMC psychiatry, 14(1), 1.

24. Murch, W. S., & Clark, L. (2016). Games in the brain: neural substrates of gambling addiction. The Neuroscientist, 22(5), 534–545.

25. Nimmrich, V., Draguhn, A., & Axmacher, N. (2015). Neuronal network oscillations in neurodegenerative diseases. Neuromolecular medicine, 17(3), 270–284.

26. Tang, Y. Y., Ma, Y., Fan, Y., Feng, H., Wang, J., Feng, S., & Zhang, Y. (2009). Central and autonomic nervous system interaction is altered by short-term meditation. Proceedings of the national Academy of Sciences, 106(22), 8865–8870.

27. Torres-García, A. A., Reyes-García, C. A., Villaseñor-Pineda, L., & Ramírez-Cortés, J. M. (2013). Análisis de señales electroencefalográficas para la clasificación de habla imaginada. Revista mexicana de ingeniería biomédica, 34(1), 23–39.

28. VaezMousavi, M., Barry, R. J., Rushby, J., & Clarke, A. (2007). Evidence for differentiation of arousal and activation in normal adults.

29. Wang, Z., & Goonewardene, L. A. (2004). The use of MIXED models in the analysis of animal experiments with repeated measures data. Canadian Journal of Animal Science, 84(1), 1–12.

